# Bacterial aerobic methane cycling by the marine sponge-associated microbiome

**DOI:** 10.1101/2022.10.21.513280

**Authors:** Gustavo A. Ramírez, Rinat Bar-Shalom, Andrea Furlan, Roberto Romeo, Michelle Gavagnin, Gianluca Calabrese, Arkadiy I. Garber, Laura Steindler

## Abstract

**Background:** Methanotrophy by the sponge-hosted microbiome has been mainly reported in the ecological context of deep-sea hydrocarbon seep niches where methane is either produced geothermically or via anaerobic methanogenic archaea inhabiting the sulfate-depleted sediments. However, methane oxidizing bacteria from the candidate phylum Binatota have recently been described and shown to be present in oxic shallow-water marine sponges, where sources of methane remain undescribed.

**Results:** Here, using an integrative -*omics* approach, we provide evidence for sponge-hosted bacterial methanogenesis occurring in fully oxygenated shallow water habitats. Specifically, we suggest methanogenesis occurs *via* at least two independent pathways involving methylamine and methylphosphonate transformations that, concomitantly to aerobic methane production, generate bioavailable nitrogen and phosphate, respectively. Methylphosphonate may be sourced from seawater continuously filtered by the sponge host. Methylamines may also be externally sourced or, alternatively, generated by a multi-step metabolic process where carnitine, derived from sponge cell debris, is transformed to methylamine by different sponge-hosted microbial lineages. Finally, methanotrophs specialized in pigment production, affiliated to the phylum Binatota, may provide a photoprotective function, closing a previously undescribed C_1_-metabolic loop that involves both the sponge host and specific members of the associated microbial community.

**Conclusion:** Given the global distribution of this ancient animal lineage and their remarkable water filtration activity, sponge hosted methane cycling may affect methane supersaturation in oxic coastal environments. Depending on the net balance between methanogenesis and methanotrophy, sponges may serve as marine sources or sinks of this potent greenhouse gas.

## Background

Sponges, globally dispersed sessile metazoans, host vastly diverse microbiomes [1] in a complex association collectively recognized as the sponge “holobiont” [2]. The pre-Cambrian fossil record [3, 4] and recent phylogenetic analyses [5] suggest that the sponge holobiont represents the most ancient of extant metazoan-microbe interactions [6]. The high-volume filter feeding of marine sponges [7] links benthic and pelagic biogeochemical processes and influences major nutrient cycles in diverse ocean ecosystems [8-13]. High microbial abundance (HMA) sponges may derive up to 35% of their mass from microbial symbionts [14, 15], accounting for a 1000-fold higher microbial load than the surrounding seawater [16]. The sponge microbiome is also responsible for producing, largely unclassified, secondary metabolites of biotechnological interest [17].

Intriguingly, sponge species seem to share a small core microbiome while hosting a large species-specific community [1]. Eukaryote-like repeat proteins, CRISPR-Cas systems, and DNA phosphorothioation are important mediators of symbiont-host recognition and defense against non-symbionts and viruses [18-22]. Some Thaumarchaeota are keystone symbionts that oxidize host-excreted ammonia [23-25], thereby, preventing toxic effects of this exudate on the sponge host [26]. Physiological studies have addressed the cycling of carbon [8, 27] and nutrients [nitrogen [24, 28, 29], phosphorus [30, 31], silicon [32]] by the sponge holobiont using both indirect and direct [33, 34] techniques, resulting in a better understanding of the holobiont functioning, its nutrient budgets and the related ecological impact [reviewed in[35, 36]]. Stable isotope incubations combined with imaging techniques (Nano-SIMS) have revealed that not only microbial symbionts but also host choanocyte sponge cells can directly take up DOM, and that it is the latter host cells that first assimilate DOM from filtered seawater, likely via pinocytosis, and then translocate the processed DOM to the symbionts inhabiting the sponge mesohyl [37, 38]. Recent efforts have characterized the community-level functional potential of sponge symbionts [18, 39] and significantly increased the number of available sponge-associated microbial genomes [40]. Genome-centered metatranscriptomic sponge surveys have explored host-microbe interactions, energy and carbon metabolism [41, 42], nitrogen cycling [24, 43], and, recently, the Tethybacterales: an uncultured clade within the Gammaproteobacteria [44]. Despite these advances, few studies have linked sponge biogeochemical cycling activities to the corresponding microbial lineages or described how these activities may influence, over geological time scales, the biogeochemical state of the planet [45]. For example, the extent to which sponges are involved in the cycling of short-chain alkanes is underexplored.

The co-detection of archaeal methanogens and sulfate reducing bacteria (SRBs) long ago suggested the presence of anaerobic niches in demosponge tissue [46]. Later work showed that sponge pumping dynamics [47] as well as distinct oxygen removal patterns not related to water pumping activity [48] elicit anoxic microenvironments where active anaerobic microbial activities such as sulfate reduction [49], denitrification, and fermentation [50] occur. Hydrocarbon degradation by sponge symbionts in deep-sea seep environments has previously been characterized and involves methylotrophic and short-chain alkane (methane, ethane, butane) specialized symbioses [51-54]. This body of work indicates that canonical archaeal methanogenesis, in anoxic tissue microniches, and short-chain alkane oxidation, in hydrocarbon rich deep-sea hydrothermal vent and cold seep environments, are activities present in sponge-associated microbiomes. Interestingly, methane oxidation activity is also predicted for sponge-associated members of the proposed Candidate Phylum Binatota [55] (also annotated as Desulfobacterota) hosted by *Petrosia ficiformis*, a sponge inhabiting fully-oxygenated shallow seas where no known hydrothermal venting or hydrocarbon seepage exists [42]. The presence of Binatota (described as Deltaproteobacteria bin18) was also reported in the shallow water growing sponge species *Aplysina aerophoba* [39]. The source of methane for methanotrophs hosted by these oxic shallow-water marine sponges remains unknown.

Here, activities of the *A. aerophoba-hosted* microbiome, residing in fully oxic seas where no hydrocarbon seepage or hydrothermal venting is known, was explored using genome-centered metatranscriptomics, gene-targeted sequencing, and metabolomics. We find no evidence for canonical archaeal methanogenesis and show that aerobic bacterial methane synthesis may occur via two recently described metabolisms: i) methylphosphonate (MPN) degradation, the proposed solution to the ocean’s methane paradox [56, 57], and ii) a recently described methylamine (MeA) [58] transformation catalyzed by a 5’ pyridoxal-phosphate-dependent aspartate aminotransferase [59]. Further, we report that microbial community processing of cell-debris generates carnitine. Carnitine may be subsequently metabolized to MeA and ultimately methane. We highlight that a potential fate of this biogenic methane, produced endogenously through either the MeA or MPN pathways, is oxidation by methylotrophic members of the Candidate Phylum Binatota, a lineage specialized in the production of photoprotective pigments that may benefit the host and, thereby, describe a novel methane-centered “metabolite processing loop” of potential symbiotic importance.

## Methods

### Sampling and sample preservation

Mediterranean *Aplysina aerophoba* specimens were sampled from the Northern Adriatic in the Gulf of Trieste (45°36.376, 13°43.1874), using SCUBA, within meters of each other. Four individuals sponge specimens were separately sampled twice in a 24h period (at 12:00 noon day 1 and 12:00 noon day 2). Immediately upon collection all tissues were *in situ* preserved in RNA Later solution using an underwater chamber as detailed elsewhere [60], and, once out of the water, kept on ice for a few hours prior to freezing and transport to shore-based storage at −80°C.

### Preparation of libraries and metatranscriptomic sequencing

RNA was extracted using Allprep DNA/RNA mini kit (Qiagen, Germany). Briefly, each extraction was preformed using 30 mg of sponge sample placed in a Lysing Matrix E tube (MP Biomedicals, Santa Ana, CA) to which RLT buffer containing Reagent DX (Qiagen, Hilden, Germany) was added. Cells were disrupted using a TissueLyser II system (Qiagen, Germany) for 30 sec at 30 Hz followed by 10 min centrifugation at maximum speed. All subsequent RNA extraction steps were performed according to the manufacturer’s protocol. SUPERase In (Life Technologies, USA) and TURBO DNA-free kit (Thermo Fisher Scientific, USA) were used for RNase inhibition and DNase treatments, respectively. RNA cleanup and concentration were done using RNeasy MiniElute kit (Qiagen, Germany). In order to achieve sufficient coverage of informative nonribosomal transcripts, rRNA was removed with RiboMinus Eukaryote System V2 kit (Ambion, Life Technologies, USA) with eukaryotic mouse-rat-human probes coupled with prokaryotic probes. ERCC RNA Spike-In Control mixes (Life Technologies, USA) were added to 5 μg of total RNA. RNA concentrations were measured using a Qubit 2.0 Fluorometer and RNA reagents (Thermo Fisher Scientific, USA), before and after rRNA depletion. In parallel, RNA integrity and purity were determined using a TapeStation 2200 system, applying the High sensitivity RNA Screen Tape assay (Agilent Technologies, USA), before and after rRNA depletion as well. Ultimately, 13ng of rRNA-depleted RNA were processed for cDNA libraries preparations using the Collibri stranded RNA library prep kit (Thermo Fisher Scientific, USA) according to the manufacturer’s protocol with the sole exception being that, following addition of the index codes, cDNA amplification was performed with 8 rather than the 9-11 recommended PCR cycles. The number of PCR cycles was optimized for our samples to reduce PCR bias. The libraries were quantified using Invitrogen Collibri Library Quantification Kit (Invitrogen, Thermo Fisher Scientific) according to the manufacturer’s guide using Real-Time qPCR. For pre-sequencing quality control (QC), 2μl aliquots of each provided library were pooled. The resulting QC pool was size and concentration checked on an Agilent D1000 TapeStation system and a Qubit 2.0 fluorometer, respectively. The pool was adjusted to 1nM and loaded on an Illumina MiniSeq Mid Output flow cell at 1.5pM. After demultiplexing the percent of each library was used to calculate new volumes to use for constructing a normalized sequencing pool. This pool was also size and concentration checked, as described above, and subsequently normalized to 2nM. The normalized pool was run on an Illumina NextSeq High Output flowcell at 2.2pM.

### Metatranscriptomic sequence processing

Paired-end Illumina libraries were inspected for quality parameters and repetitive sequences using the FastQC software package. Adapter trimming was performed using the trim adapters bbduk script from BBMaps (https://sourceforge.net/projects/bbmap/). Trimmed paired-end files were interleaved for alignment against rRNA libraries using SortMeRNA [61]. Non-aligned reads were subsequently split into paired forward and reverse files for downstream analyses. A *de novo* co-assembly was performed using merged forward and reversed adapter trimmed and non-rRNA aligned sequences with rnaSPAdes v.3.14.1 [62]. Sequence counts at each step for all libraries, in addition to co-assembly summary statistics, are provided in Table S1.

### 16S rRNA gene and transcript ASV analysis

As detailed above, RNA and DNA were extracted in parallel and RNA was subsequently reverse transcribed. DNA and cDNA extracts were used as templates for PCR-based amplification for 16S rRNA gene/transcript V4 amplicon generation using the following primers: 515F-ACACTGACGACATGGTTCTACAGTGCCAGCMGCCGCGGTAA and 806R-TACGGTAGCAGAGACTTGGTCTGGACTACNVGGGTWTCTAAT, and thermocycling program: 94°C for 3 mins, followed for 32 × [94°C for 45s, 50°C for 60s, 72°C for 90s], 72°C for 10 mins, and a 4°C hold. Amplicon sequencing was performed and an Illumina MiSeq platform. Sequence analyses were performed using the DADA2 package [63] implemented in R. Briefly, forward and reverse reads were trimmed with the AlterAndTrim() command using the following parameters: trimLeft = c(20,20), maxEE = c(2,2), phix = TRUE, multithread = TRUE, minLen = 120, followed by error assessments and independent forward and reverse read dereplication. Sequencing errors were removed using the dada() command and error-free forward and reverse reads were merged using the mergePairs() command, specifying overhand trimming and a minimum overlap of 120 base pairs. The resulting amplicon sequence variants (ASVs) were assigned taxonomy by alignment against the SILVA 132 database[64]. ASV count tables and taxonomy assignments were merged into an S4 object for diversity analysis and summary visualization using vegan in a phyloseq[65].

### Pathway completion estimates

Prodigal v2.6.1 [66] predicted protein products were annotated against the KEGG database [67] via GhostKOALA [68] with the following parameters: taxonomy group, Prokaryotes; database, genus_prokaryotes + family_eukaryotes; accessed February 2021. The output annotation file was used for pathway completion assessment and visualization using KEGG-decoder.py [69].

### Read mapping

To assess coverage as a proxy for transcript abundance, quality-trimmed non-rRNA short reads were mapped to our *de novo* metatranscriptomic assembly and a public reference set of MAGs binned from *A. aerophoba* in close geographical proximity to our study site [39] using Bowtie2 [58] and the following parameters: read counts were normalized to Transcripts per Million (TPM) per library, as suggested elsewhere [70], and all data was concatenated into read count tables for downstream statistical analyses.

### Phylogenomic tree

A phylogenomic tree was generated for *A. aerophoba* derived MAGs using the GToTree package [71]. Briefly, 37 publicly available MAGs [39] were used as the input and were ran against a GToTree’s “Bacteria” HMM collection of single copy genes within this domain resulting in a concatenated protein alignment constructed using Muscle [72] and trimmed with TrimAl [73]. The tree was constructed in FastTree2 [74] and visualized using FigTree (https://github.com/rambaut/figtree).

### ORF calling, annotation, and targeted gene analyses

Open reading frame (ORF) identification and, subsequently, prokaryote predicted protein product annotations were performed with Prodigal v2.6.1 [66] implemented in Prokka [75]. Selected ORFs were also aligned against the NCBI non-redundant (nr) database (accessed in April 2021) using BLASTp for closest homologue taxonomy and functional annotation supplementation. Targeted single gene homologue searches within our data were also performed using BLASTp 2.2.30+ (*E*-value threshold = 1E–30, identity = 50%) against predicted protein sequences inferred from i) metatranscriptomic assemblies, ii) unbinned metagenomic contigs, and iii) MAGs.

### Metabolome analysis

Four ~1.5g frozen tissue samples collected from specimens 25, 27, 28, and 29 each in 1:1 wt:wt sample to EtOH ratio solutions were shipped on dry ice for biogenic amines panel metabolome analyses by HILIC-QTOF MS/MS [76] at the UC Davis West Coast Metabolomic Center (http://metabolomics.ucdavis.edu/). Raw output files were curated based on internal standard removal, signal to noise ratio cutoff and a minimum peak threshold of 1000, and subsequently parsed based on average (*n*=4) metabolite log fold increases relative to blanks, as suggested elsewhere [77]. Only metabolites with non-redundant names and InChIKey identifiers and average fold changes higher than 5 relative to blanks were retained for further analyses.

## Results and Discussion

### Net relative activity of symbiont lineages

A normalized activity survey based on metatranscriptomic read mapping against 37 Metagenome Assembled Genomes (MAGs) [39] representative of the *A. aerophoba-associated* microbial community (Figure S1) shows Gammaproteobacteria, Cyanobacteria, Deltaproteobacteria, Acidobacteria, Chloroflexi, and Poribacteria as the most active lineages (Figure 1, Figure S2). The Alphaproteobacteria appear to be an abundant and diverse lineage with relatively lower levels of activity. Our observations corroborate previous work regarding sponge microbiome activity [43] and highlight consistent transcriptional activity, *i.e.*, relative activity of each community lineage is constant across time (24h) and different sponge specimens (*n* =4, Figure S2).

**Figure 1.**
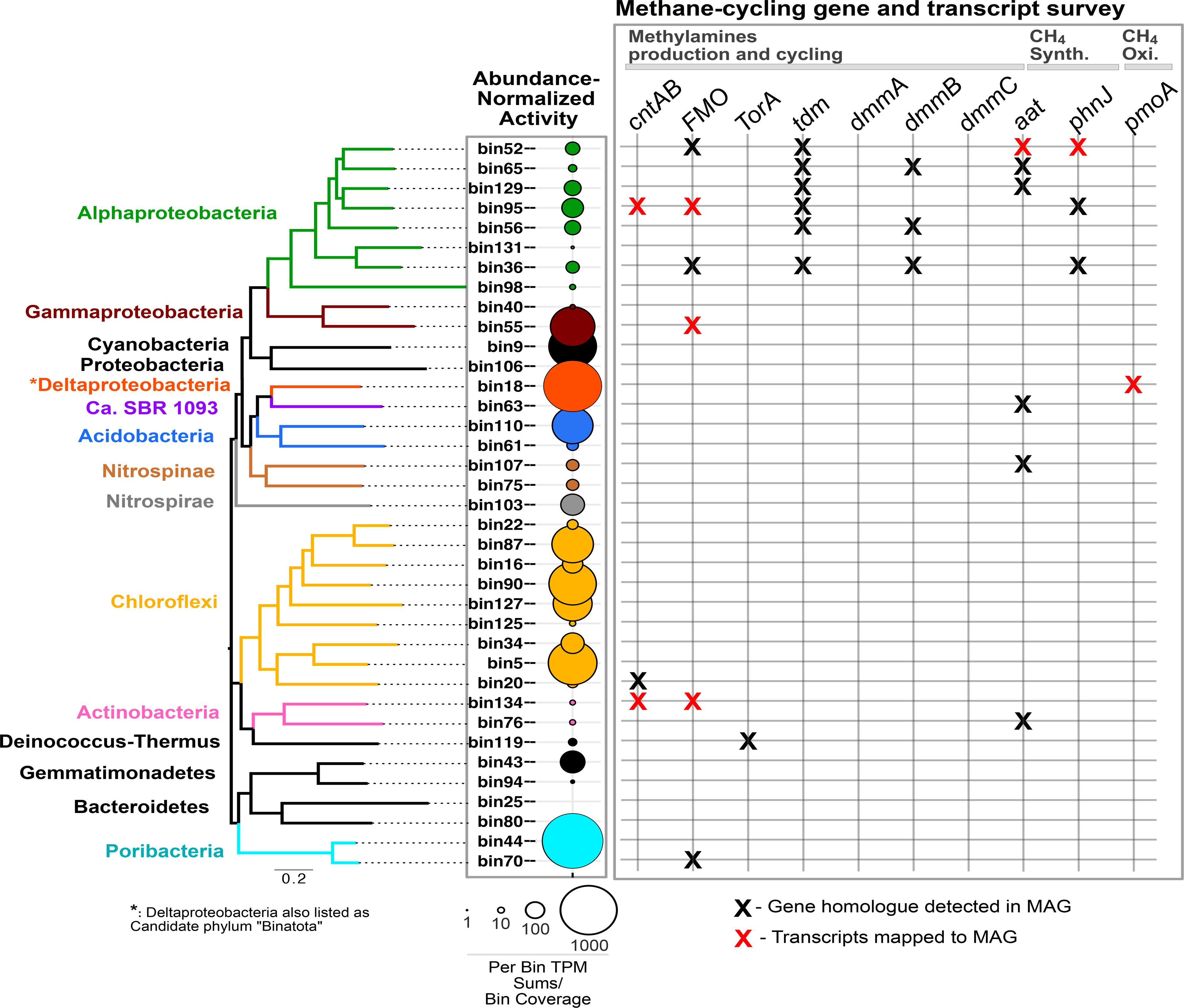
Survey of methane cycling genes and transcripts. Left - Relative global activity survey for 37 dominant lineages, arranged based on phylogenomic relatedness and color-coded at the Phylum-level of taxonomy, in the *A. aerophoba*-associated microbiome. Relative activity is represented by circle size and depicts mean values per lineage from eight independent metatranscriptomes: four sponge specimens sampled twice, at noon twenty-four hours apart (see Figure S2 for individual results). Right - Methane relevant genes and gene transcripts detected in MAGs and in the metatranscriptomes are depicted by black and red “X” symbols, respectively.

### Sponge microbiome predicted activities: C-, N-, and S-cycling, fermentations, and photosystems

As a broad metatranscriptomic-predicted activity survey of the *A. aerophoba*-associated microbiota, aerobic and anaerobic metabolic pathways [49] were explored (Fig 2 & S3). Carbon fixation via oxygenic photoautotrophy (CBB cycle) predominates. The incomplete transcription of other potential chemoautotrophic pathways [e.g.: 3-Hydroxypropionate (3HP) Bicycle, and 4-Hydroxybutyrate/3-Hydroxypropionate (4HB/3HP)] is also observed. Previous transcriptomic and metatranscriptomic surveys report the presence of CBB and rTCA cycles in Demosponge associated symbionts [41, 78]. Here, we corroborate an active CBB cycle, but fail to detect transcripts associated with the rTCA pathway suggesting that the rTCA cycle may not be a ubiquitous feature across sponge microbiomes.

**Figure 2.**
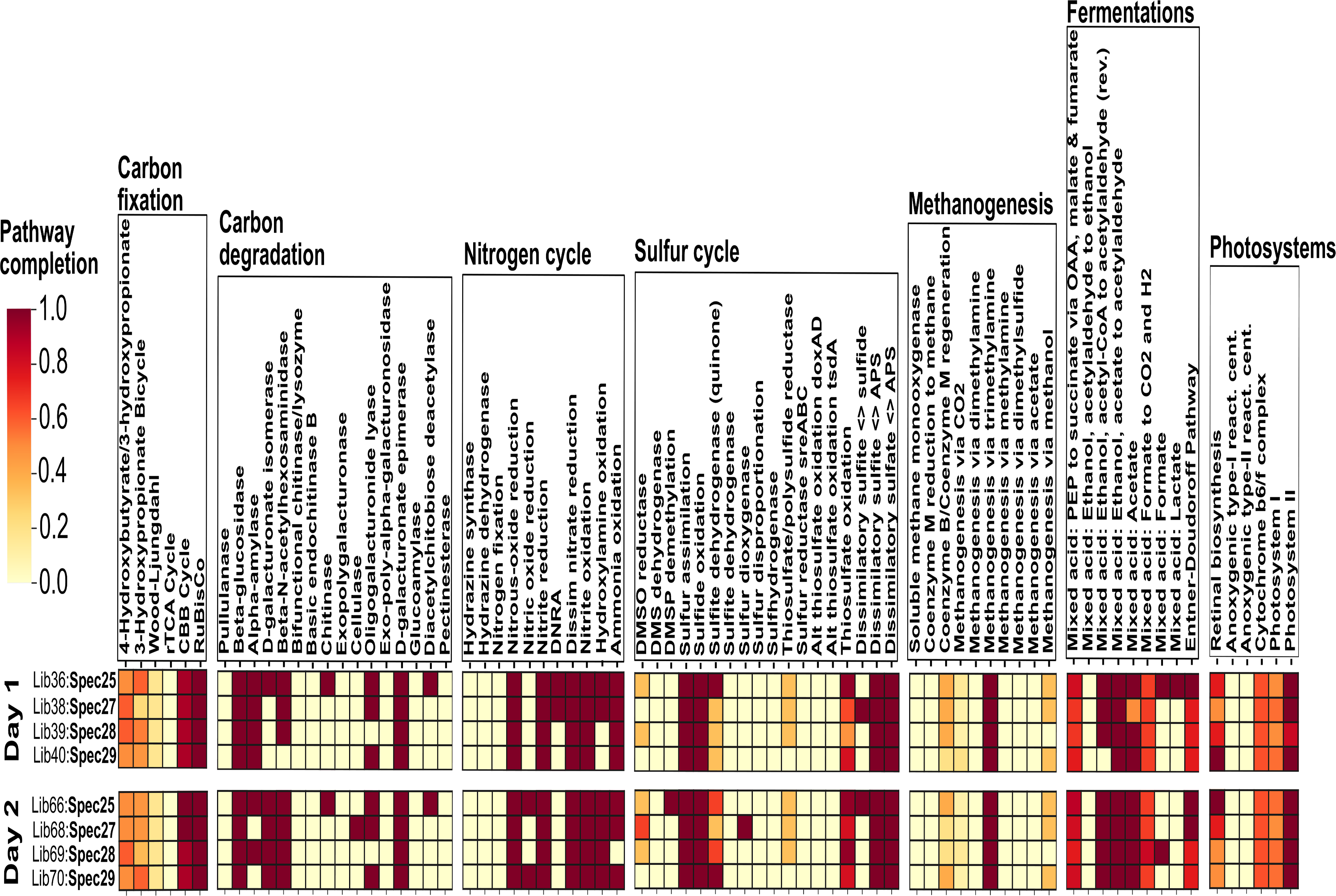
Microbiome metatranscriptome-predicted activity. Microbial activity survey based on collection day and specimen source for selected microbial metabolisms: carbon fixation, carbon degradation, nitrogen cycle, sulfur cycle, methanogenesis, fermentation, and photosystems. Day 1 and Day 2 depictions represent biological replicates collected 24h apart at noon. The color gradient depicts the fractional percentage of KEGG module pathway completion.

Active carbon degradation pathways include those of pectin, chitin, cellulose, (D-galacturonate epimerase, D-galacturonate isomerase, oligogalacturonide lyase, chitinase, cellulase), and alphaamylase utilization. The catalysis of plant cell walls (pectin and cellulose) and continuously regenerated sponge matrix components (chitin) posits heterotrophy as a functional motif for this symbiosis, as previously suggested [79]. Interestingly, diacetylchitobiose deacetylase activity is a chitin degradation pathway only found in Archaea [80] and shows the heterotrophic activities of these community members (Figure S3).

An active sulfur cycle is predicted in our samples (Figure 2), adding to a growing body of evidence showing that microbial sulfur cycling is an important sponge resource [81-83]. Continuous sulfur cycling, localized to anoxic niches within sponges, may benefit the host by the removal of toxic metabolites such as hydrogen sulfide [49, 84]. Diversification of sulfur metabolism in bacteria coincides with the emergence of metazoan life [85], suggesting a long co-evolutionary history reflected in the reduced genomes, as reviewed elsewhere [86], and versatile metabolisms of contemporary sulfur cycling sponge symbionts [82, 83].

Transcripts related to acetate and ethanol-based mixed acid fermentations are detected and suggest activities with reactions obligatorily localized to anaerobic niches [49]. Complete pathways for thiamin, riboflavin, and cobalamin synthesis *(vitamins* B_1_, B_2_, and B_12_, respectively) and for all 20 essential amino acids and B vitamins, important microbial products with potential host benefits, are expressed across all biological replicates (Figure S3) and support metabolite exchange [87] as one potential driver for this symbiosis.

Photoactivity is dominated by cyanobacterial Photosystem II transcripts and corroborates previous reports [60, 88]. Photosynthate transport from symbiotic photoautotrophs to the sponge host has been suggested as a common activity underpinning this symbiosis [60, 88, 89] but the transfer of photosynthates to the sponge is host- and cyanobacteria-specific and cannot be predicted by -*omics* data alone [42].

Nitrogen cycling activity was also expressed and is described and discussed later in relation to the here suggested methane production pathways.

Interestingly, we detected a transcribed predicted pathway for methanogenesis via trimethylamine (TMA) (Figure 2), a known activity of archaea in the human gastrointestinal tract [90].

### *Canonical anaerobic archaeal methanogenesis is absent in* A. aerophoba

Motivated by the presence of methanotrophs (Binatota) and by the unexpected KEGG-pathway prediction for TMA-based methanogenesis (Figure 2), we used a combined -*omics* approach to explore methane production avenues in *A. aerophoba.* We do not detect archaeal methanogenic lineages canonically associated with this activity (Figure S1) nor genes or transcripts for euryarchaeal methyl-coenzyme M reductase (*mcrA*). Additional scrutiny of transcripts encoding TMA methyltransferase homologues *(mttB* within the COG5598 super family), revealed that these genes lack pyrrolysine (Figure S4), a characteristic non-canonical amino acid residue involved in methane cleavage from methylated amines [91]. Non-pyrrolysine *mttB* homologues allow the strict anaerobe *Desulfitobacterium hafniense* to grow on glycine betaine with carbon dioxide and dimethylglycine as by-products [92]. Accordingly, we hypothesize that these *mttB* homologues may still play an important role in TMA cycling, however, the functional prediction of TMA methylotrophic methanogenesis by Euryarchaeal *mttB* is likely not correct, and alternative methane sources for Binatota methanotrophy in *A. aerophoba* were investigated.

### Sponge-hosted aerobic bacterial methane synthesis: marine methane paradox

Studies of the oversaturation of methane in fully oxygenated aquatic environments, a phenomenon dubbed the “marine methane paradox” [93] in the ocean, have revealed that aerobic bacterial degradation of methylphosphonates [56] and methylamines [59] results in methane production. These observations challenge the view of methanogenesis as a strictly anaerobic process performed exclusively by archaea and show that i) bacteria play an important role in the production of a potent greenhouse gas and that ii) this activity occurs in a broader range of redox (*e.g*., aerobic, microaerophilic) and chemical (sulfidic) environments. We, thus, searched for the presence and expression of marker genes for these aerobic methanogenesis pathways in sponge-derived MAGs [39] and our metatranscriptomes.

### Methylphosphonate-based aerobic bacterial methane synthesis

An alternative pathway for methane generation is the aerobic degradation of methylphosphonates (MPNs) through the C-P lyase activity of PhnJ [56]. This pathway is enriched in free-living marine pelagic bacteria from phosphate limited locations including the Sargasso and Mediterranean Seas [94], the latter being the sampling site of our sponge specimens. Genes encoding *phnJ* homologues are transcribed across all our metatranscriptomic libraries (Figure 3). We also detect *phnJ* gene homologues in 3 Alphaproteobacterial MAGs, with active transcription of this gene detected in at least one lineage (bin52, Figure 1). Phosphonate metabolism was previously reported for the sponge *Xestospongia muta* [81] which, together with our results, suggests that seawater derived phosphonates may be an important source of inorganic phosphate for sponge-associated microbes. An additional mechanism for phosphate sequestration in phosphate-limited conditions is the production of polyphosphates, which was previously reported for sponge symbionts [13]. Overall, PhnJ activity suggests that sponge symbionts may experience phosphate-limited conditions and that mechanisms to cope with this nutrient limitation are important for symbiont survival. We highlight that these results also imply that hydrocarbons, including methane, are potentially released as a result of phosphonate cleavage by C-P lyase.

**Figure 3.**
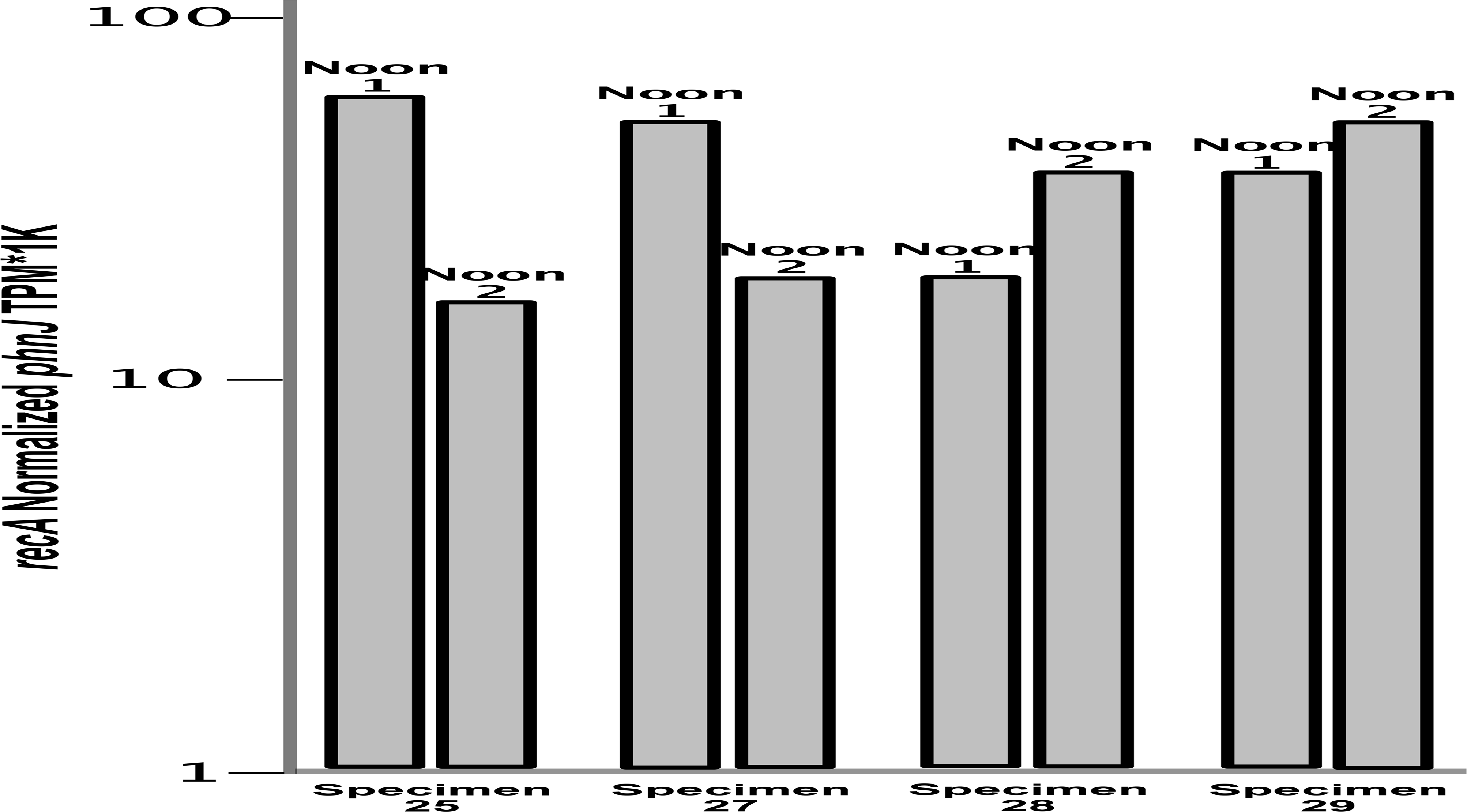
*phnJ* expression of a sponge-hosted Alphaproteobacteria. *recA* normalized transcriptional activity of the *phnJ* gene in four sponge specimens (samples 25, 27, 28, 29) and in two time replicates (Noon 1 and Noon 2).

### Methylamine-based aerobic bacterial methane synthesis

Recently, a new MeA-based aerobic methane synthesis metabolism, using 5’ pyridoxalphosphate-dependent aspartate aminotransferase, was described in freshwater bacteria [59]. We report the transcription in our metatranscriptomic libraries of closely related gene homologues to the functionally confirmed proteobacterial *aat* (sequence: MK170382.1 recovered from *Acidovorax* sp.), the single heritable unit required to confer methane generating activity to an *E. coli* clone (Figure 4). The closest transcribed homologues to MK170382.1, classified as Alphaproteobacterial predicted proteins, also maintain conserved key functional domains: a catalytic Lys residue and nine pyridoxal 5’-phosphate biding sites (Figure 4 and S5). Additionally, *aat* gene homologues detected in MAGs allowed assignment of Alphaproteobacteria, Actinobacteria, Nitrospinae, and SBR phylogenetic provenance. At least one of these linages (Alphaproteobacterial bin52) actively transcribes this gene (Figures 1 & 4). Noting that AAT expressed in *E. coli* confers aerobic methane synthesis ability in the presence of MeA [59], our genomic and metatranscriptomic findings suggest that demosponges may host aerobic bacterial methane synthesis *via* AAT-mediated MeA metabolism. Methane generation via *aat* expression is also linked to growth in *Acidovorax* and when heterologously expressed in *E. coli*, suggesting that methanogenic activity may concomitantly allow anabolic nitrogen uptake. *aat*-based methane release from MeA rather than the oxidation of the MeA-methyl groups for energy may only be favored under high carbon and low nitrogen conditions. Whilst sponge-secreted ammonia may result in bioavailable nitrogen to the associated microbiota, conditions of nitrogen availability inside the sponge are variable. For example, the fluxes of dissolved inorganic nitrogen differed across specimens of *X. muta*, with some specimens serving as source and others as sinks of dissolved inorganic nitrogen [87, 95]. Furthermore, *A. aerophoba*, was shown to serve as ammonium sink in spring and as ammonium source in the fall [24]. Some sponge species are characterized by constitutive nitrogen fixation [96] which may provide continuous availability of inorganic nitrogen to the microbial community, however, in the case of *A. aerophoba*, we could not detect genes involved in nitrogen fixation based on our metatranscriptomics predictions (Figure 2). Finally, whilst our metatranscriptomic analysis predicts ammonia oxidation coupled with nitrite oxidation, as well as dissimilatory nitrate reduction (i.e., nitrate ammonification), it also predicts denitrification, which results in loss of bioavailable nitrogen sources to the microbiota (Figure 2). Denitrification was previously shown to occur in the Mediterranean sponge species *Dysidea avara* and *Chondrosia reniformis* [97]. Denitrification can be favored over other nitrogen cycling pathways in anaerobic niches within the sponge tissue, possibly resulting in nitrogen limited conditions to the microbiota. Under these conditions MeA may serve as a nitrogen source to part of the microbial community and the process would result in methane release. Future work involving functional characterization of sponge-hosted *aat* homologues is needed to test our proposed involvement of *aat* in sponge-associated aerobic methane synthesis.

**Figure 4.**
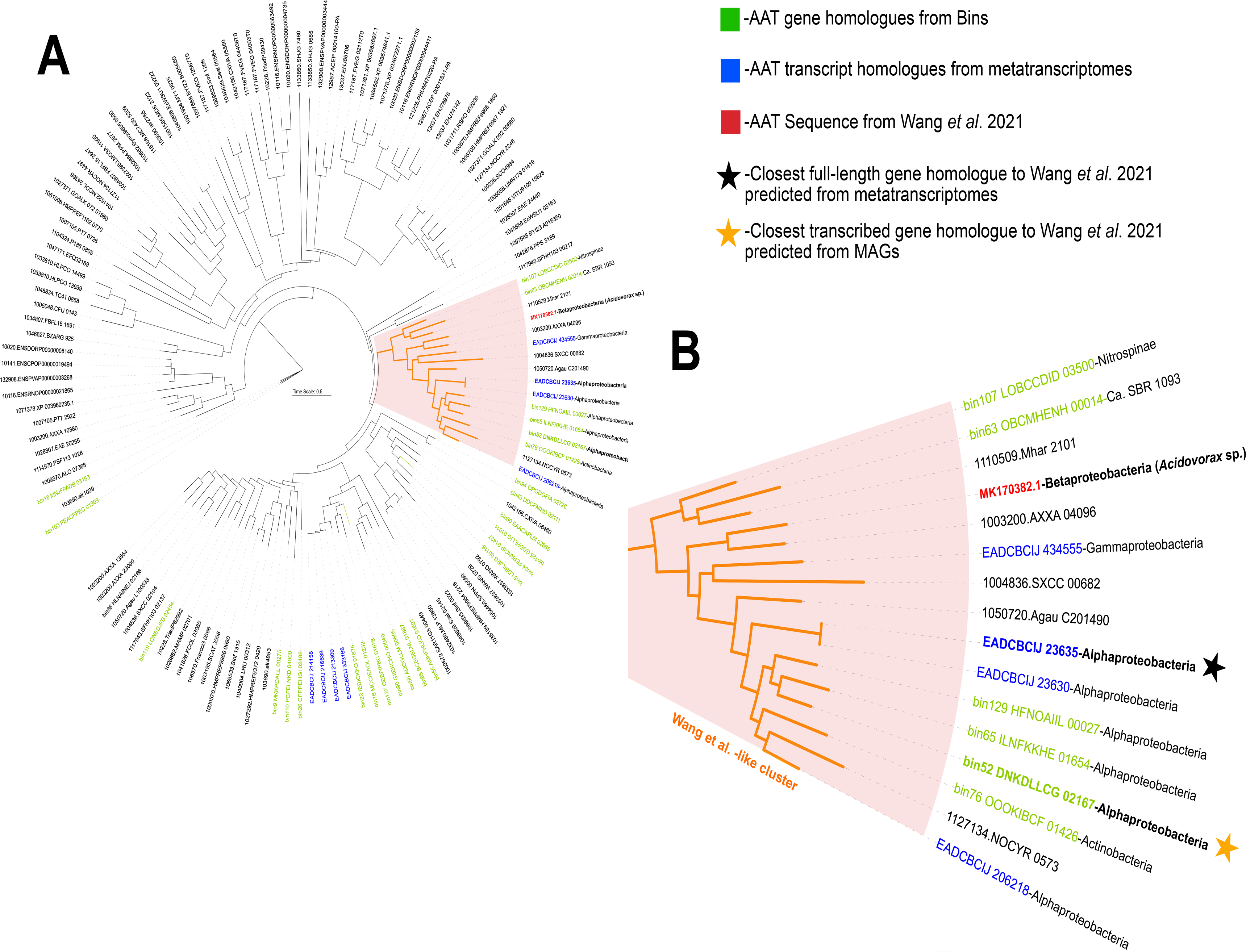
Phylogeny of *aat* genes and transcripts. (A) Phylogenetic tree of 100 COG0446 member sequences including *aat* gene (green leaves) and transcript (blue leaves) homologues identified from *A. aerophoba* MAGs and metatranscriptomes, respectively. The closest sponge-recovered gene and transcript sequences to MK170382.1 (red leaf), the *aat* gene confirmed to confer the methanogenesis phenotype in *E. coli* by Wang and colleagues [59], are emphasized in the orange cluster. (B) A focused depiction of the Wang and colleagues cluster highlighting the phylogenetic provenance of all highly similar sequences to MK170382.1 and showing the closest full-length genes recovered from MAGs and metatranscriptomic assemblies with gold and black stars, respectively.

### Methylphosphonate and methylamine sources in marine sponges

The source of MPN and MeA may be exogenous, with the sponge concentrating these compounds from DOM during its efficient water filtration. MPN and MeA have in fact previously been reported as common compounds in seawater [56, 98–100]. Alternatively, these substrates for methanogenesis may derive from endogenous metabolism. Based on our inability to detect methylphosphonate synthase (*mpnS*) genes or transcripts in our data, encoding the enzyme responsible for MPN synthesis [101], we propose that MPN is not produced endogenously by the sponge-associated community and rather likely sourced from the surrounding seawater. Conversely, the source of MeA may indeed be endogenous, deriving from the metabolic processing of TMA [102]. TMA can be produced from choline [103], glycine betaine [104] or carnitine [105]. Genes and transcripts related to TMA generation from glycine (*grdH*) and choline (*cutC*) derived transformations were not detected in our genomes and metatranscriptomes. We, however, report the presence of gene homologues to *cntAB*, involved in carnitine-based TMA biosynthesis [105]. We performed metabolomic analysis which directly detected the presence of carnitine in our *A. aerophoba* tissue samples (Supplemental File 1). Carnitine (γ-trimethylamino-β-hydroxybutyric acid) is a ubiquitous quaternary amine produced by all members of animalia, including Porifera [106] that is metabolized by prokaryotes under aerobic conditions into TMA and malic acid or, under anaerobic conditions, to glycine betaine and, subsequently, glycine [107]. Recent studies of *A. aerophoba* used metagenomics and single cell sequencing to infer the presence of a symbiont guild specialized in carnitine utilization as: i) a source of carbon and nitrogen anabolism [39] and/or ii) a substrate for energy-yielding catabolism [18]. Further, the presence of 2-methylbutyryl-carnitine was reported for six sponge species from the Great Barrier Reef, Australia [108], supporting the ubiquitous presence of carnitine-related compounds in sponges and their potential not only as a source of food but also as precursor to methanogenesis substrates for the associated microbiota. In fact, microbial-mediated production of TMA from carnitine in our samples is predicted by the transcription of the *cntAB* genes by lineages classified as Alphaproteobacteria and Actinobacteria (Figure 1). Further, TMA oxidation to trimethylamine-N-oxide (TMAO) and “back-production” from TMAO are predicted by the detection of FMO transcripts from Proteobacteria and Actinobacteria and *torA* genes in Actinobacteria, encoding TMA oxidase [109] and TMO reductase [110], respectively. The detection of genes homologous to *tdm* and at least one subunit of the *dmmABC* complex in Alphaproteobacterial lineages, encoding TMAO demethylase [109] and DMA monooxygenase [111], respectively, also suggests that TMAO may be further oxidized to dimethylamine (DMA) and, finally, MeA (Figure 1). Taken together, we suggest that, while both substrates for aerobic bacterial methanogenesis (MPN and MeA) may be seawater derived, MeA may also be endogenously produced through the recycling of sponge cell debris (i.e., carnitine) resulting from continuous replacement of choanocyte cells. A high turnover of choanocyte cells, involving high proliferation followed by cell shedding, was shown to occur in *Halisarca caerulea* and was suggested to enable the constant renewal of the sponge filter system required for its efficient water filtration [112].

### Deltaproteobacteria (Binatota) methylotrophs as potential methane sinks

With two potential sources of biogenic aerobic methane in the sponge holobiont, the highly active *A. aerophoba*-associated symbiont bin18, (Candidate phylum Binatota, or Desulfobacterota, an unclassified Deltaproteobacteria lineage according to NCBI taxonomy) is here identified as a likely methane sink (Figures 1 and S2). A previous detailed study of this lineage described them as pigment production specialists with a predicted lifestyle that is heavily reliant on aerobic methylotrophy and alkane degradation [113]. We show that bin18 actively transcribes methane monooxygenase (*pmoA*) gene (Figure 5A). Similarly, a member of the same phylum was recently reported to express *pmoA* in the sponge species *P. ficiformis* [42] implicating this Deltaproteobacterial (Binatota) lineage in sponge methane oxidation. Methylotrophy results in reducing equivalents through the formation of methanol as a key intermediate [114], and can thus facilitate the production of carotenoid pigments. Further, we detect the transcription of genes involved in all the intermediate steps necessary for 7, 8-dihydro beta-carotene, chlorobactene, and isorenieratene pigment biosynthesis (Figure 5B). These pigments can serve as photoprotectants for the host and its symbionts [115]. Together, our results suggest that at least two independent methane production pathways, using host-derived MeA and seawater-derived MPN, represent previously unrecognized syntrophies that, through methane as an intermediate, contribute to the production of photoprotective pigments that may benefit the sponge holobiont.

**Figure 5.**
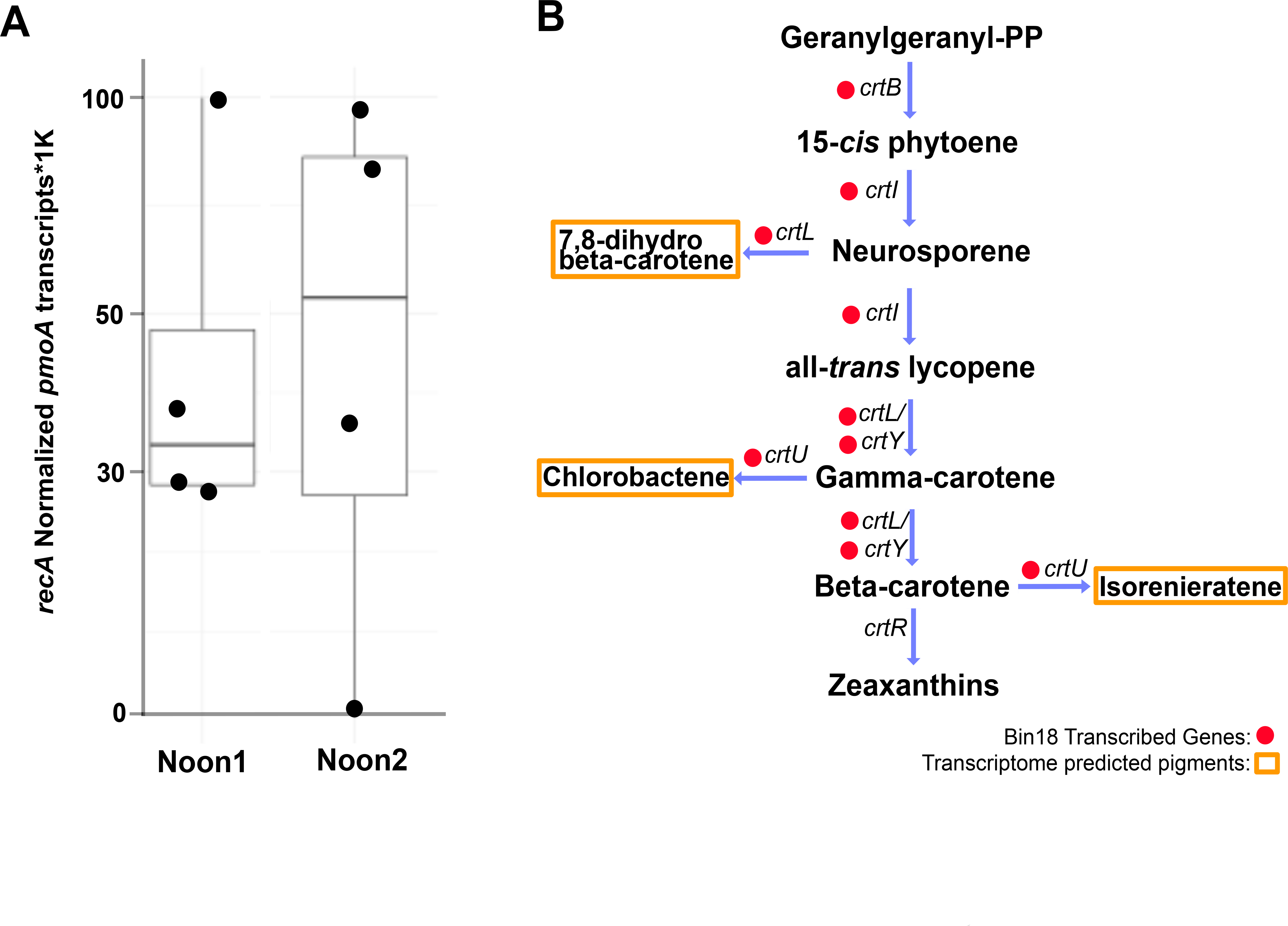
Transcription of Deltaproteobacterial (bin18, Candidate Phylum Binatota) *pmoA* gene and pigment biosynthesis pathways. (A) *recA* normalized transcriptional activity of*pmoA* gene in four sponge specimens and two time points. (B) Transcribed genes related to pigment biosynthesis pathways identified in the Deltaproteobacterial MAG are denoted by red circles next to the gene names. Predicted pigment metabolites from this activity are highlighted in yellow boxes.

### Global distribution of sponge-associated methylotrophs

To determine the global prevalence of sponge-associated methylotrophic pigment producers, we searched the Sponge Microbiome Project (SMP) dataset for highly similar (>98% ID, >99% subject length alignment) matches to the 16S rRNA gene sequence recovered from Binatota (bin18). We identify numerous matches to our Binatota query sequence in at least 46 sponge species (Figure S6) with significant (student t-test, P_val_ < 0.05) enrichment of this lineage in 15 sponge species relative to seawater metagenomes (Figure 6). The highest percent Binatota abundances in the SMP dataset are observed in the following sponge species: *Pseudocorticium jarrei, Ircinia variabilis, Cacospongia mollior, Ircinia oros*, and *Spongia agaricina*. We note that Binatota phylotypes are significantly enriched in *A. aerophoba* and *Aplysina archeri* specimens relative to seawater (Figure 6); however, this enrichment is not detected in six other *Aplysina* species sampled (Figure S6). This suggests that microbial methane cycling potential may be a species-specific activity in the genus *Aplysina*. Interestingly, *P. ficiformis*, another Mediterranean sponge species with recently published metatranscriptomes, metagenomes, and MAGs [42, 60], was also identified as hosting a significantly enriched Binatota community and was, accordingly, used as a second model for assessing methane cycling potential in sponges.

**Figure 6.**
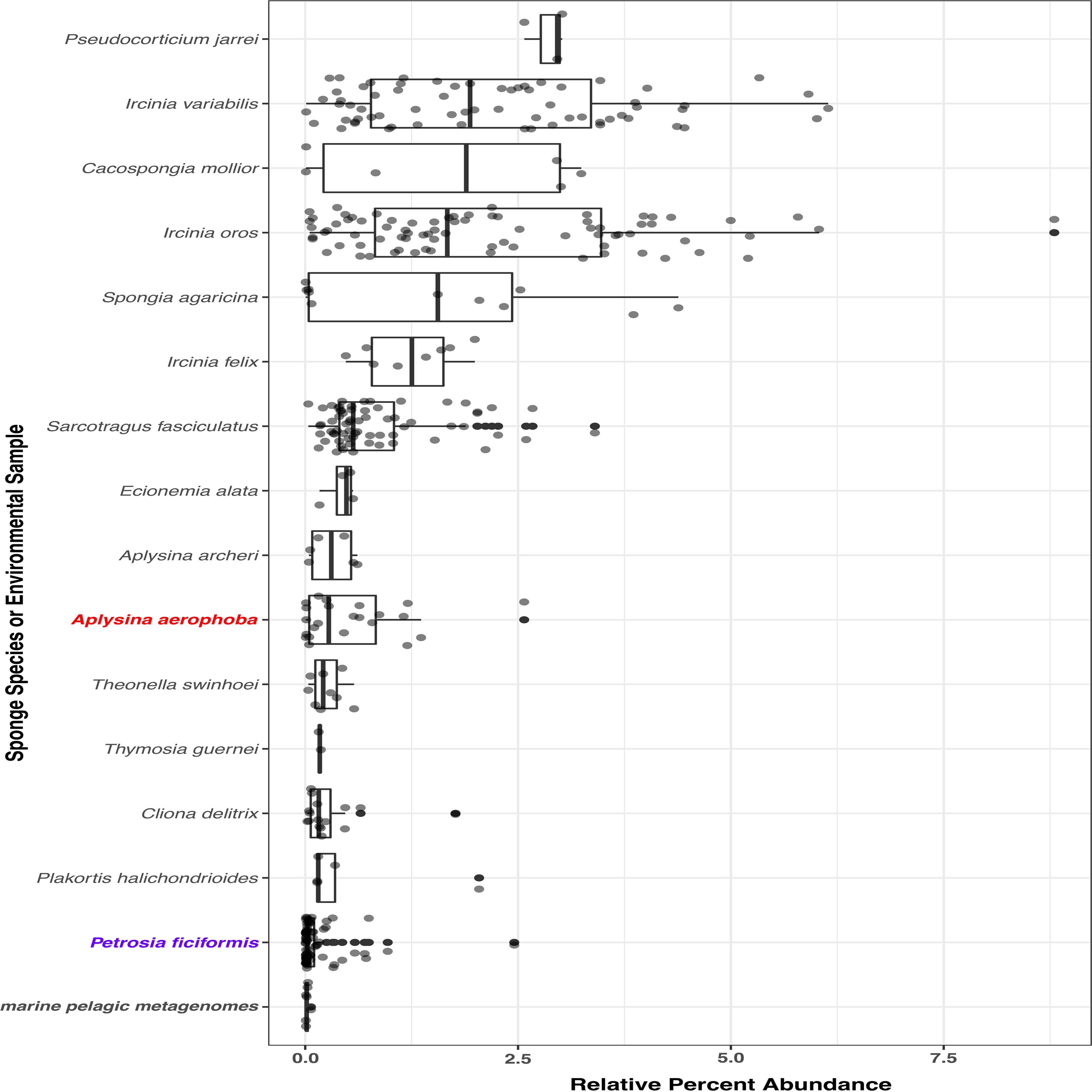
Global sponge associated Binatota survey. Relative percent abundance of 16S rRNA sequences with >98% sequence ID and 99% of query length to the *A. aerophoba* bin18 (Binatota) 16S rRNA gene sequence, summarized in quartile boxplots. Species shown had significantly (student t-test, P_val_ < 0.05) higher relative abundances of bin18 16S rRNA gene matches than marine pelagic metagenomes. Quartile box plots are arranged in descending median values of all observations from top to bottom. *Aplysina aerophoba* and *Petrosia ficiformis*, both sponges analyzed in this study, are highlighted in red and purple, respectively.

### *Methane cycling meta-analysis of* Petrosia ficiformis

To explore whether MPN- and MeA-based methane cycling activity is unique to *A. aerophoba*, or rather a more widely distributed characteristics of sponge holobionts, we re-analyzed recently published collections of MAGs and metatranscriptomes of the species *P. ficiformis* [42, 60]. Genes involved in carnitine breakdown to methylamines (*cntAB*, *tdm*, *dmmB*), MeA-based methane generation *(aat)* [59], and methane oxidation *(pmoA)* were detected in *P. ficiformis* MAGs (Figure S7). In both sponge species, MeA production potential is predicted predominantly from Alphaproteobacterial community members. Additional predicted aerobic methanogenic lineages in both sponge species include members of the Actinobacteria, while in *P. ficiformis* Bacteroides, Latescibacteria, and Chloroflexi may also be involved in this activity. The gene marker for MPN-based methanogenesis *(phnJ)*, present and actively transcribed in *A. aerophoba* symbionts (Figure 3), was not detected in *P. ficiformis* MAGs nor was it found expressed based on its metatranscriptomes. These results indicate that bioavailability of phosphate to symbionts may differ across demosponge species. Lastly, we note that *pmoA* gene homologues and transcripts, indicative of methane oxidation, were also detected in a Deltaproteobacterial MAG, classified as Candidate Phylum Binatota, from *P. ficiformis* [42]. These meta-analysis results show that methane cycling may be widespread among Porifera. While the Binatota appear to serve as a common methane sink in both *A. aerophoba* and *P. ficiformis*, the microbial phyla responsible for methanogenesis and the pathways involved, appear to be species-specific (Figures 1 & S7).

### Local and global ecological impact of methane cycling in marine sponges

Symbionts enable their animal hosts to benefit from microbial metabolism and, ultimately, impact ecosystem health and function [2]. We show one such case study involving the biogeochemical cycling of C, N, and P, via two independent pathways for aerobic bacterial methane synthesis, to produce photoprotectant pigments that directly benefit the sponge host: host derived organic matter (carnitine) + environmentally filtered compounds (*e.g.*, MPN) → *in situ* aerobic bacterial methane production → generation of bioavailable N and P for symbionts + photoprotective pigments for host (Figure 7).

**Figure 7.**
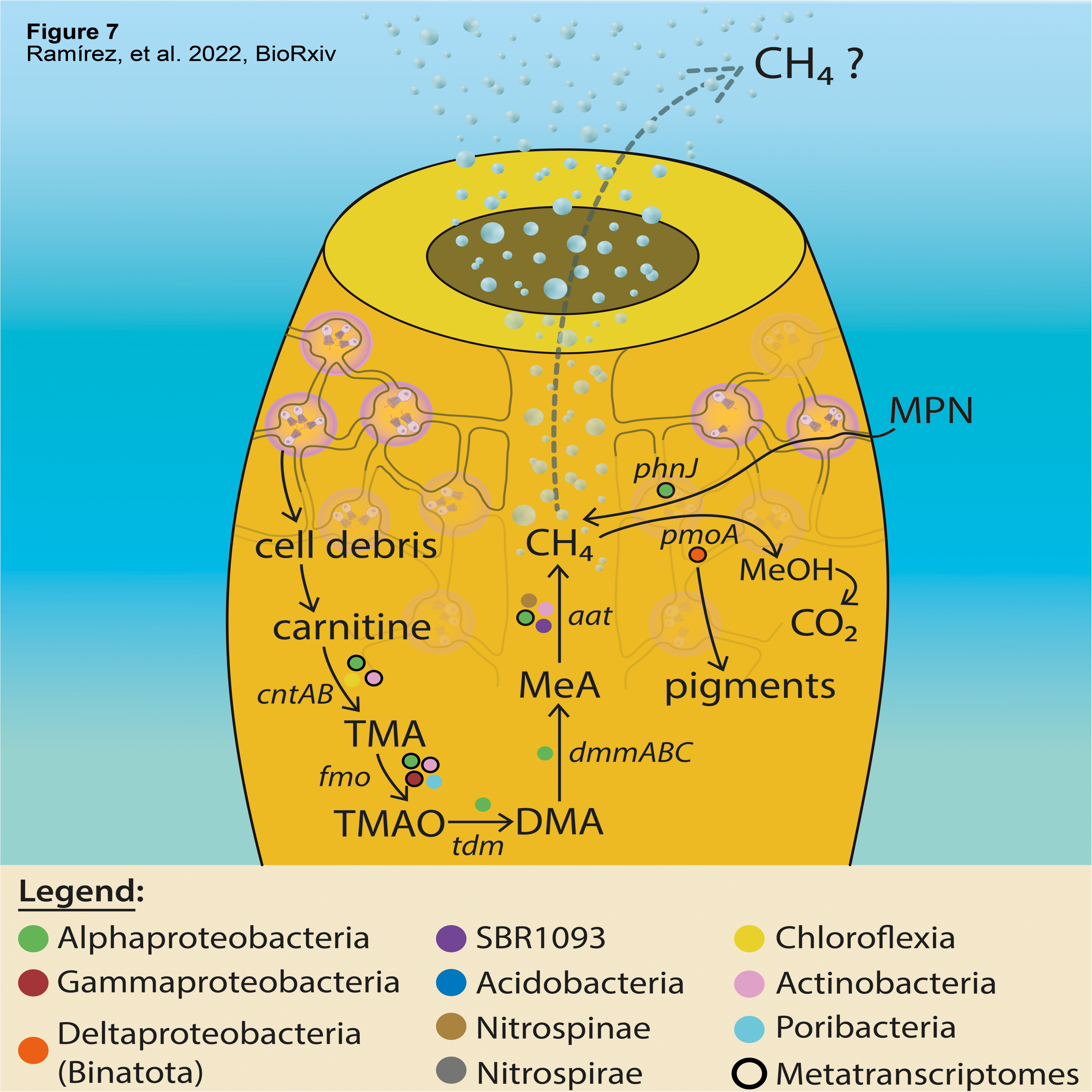
Bacterial aerobic methane cycling model. Conceptual diagram depicting a metabolic cascade involved in major microbial transformations leading to methane production/oxidation. Genes for key enzymes involved in each substrate transformation are denoted in italics. The phylogenetic association of bins where they have been detected are shown as color-coded circles next to reaction arrows. Circles with black frame also recruited transcript reads for the depicted marker gene, meaning the function was active at the time of sampling.

Methane is a potent greenhouse gas capable of trapping 3.7 times more radiated heat than CO_2_ [116] and a significant determinant, even in trace amounts, of the Earth’s radiative atmospheric balance [117]. The ocean is an important source of global methane emissions [118, 119].

Depending on the coupling of methanogenesis and methanotrophy in sponges, methane may be internally cycled, consumed, or emitted to the surrounding seawater. If emitted, considering the over half-billion-year-old history of sponge symbiosis [3, 4] and the remarkable water filtration activity of marine sponges [7], it is possible that this previously unrecognized marine animal-hosted methane cycle may have, over geological time, influenced methane concentrations of marine environments and possibly fluxes of methane from supersaturated ocean waters to the atmosphere.

## Conclusions

The metabolic activities of the sponge microbiome are saliently diverse; interestingly, here we also predict aerobic bacterial methane synthesis based on independent methylphosphonate and methylamine metabolisms. Together with the presence of abundant and active methylotrophic community members, this suggests the existence of a previously unrecognized aerobic bacterial methane cycle in demosponges that may affect methane concentration in sponge dominated marine habitats. Further studies including quantitative methane measurements in sponge incurrent and excurrent seawater, in the presence of natural and amended methanogenic substrates (MeA and MPN), and under different nutrient (N and P) conditions, will enhance our understanding of the influence of sponge holobionts on present and future marine ecosystems.

## Supporting information

Ramirez_SI_2022

## List of abbreviations

3HP: 3-Hydroxypropionate
4HB/3HP: 4-Hydroxybutyrate/3-Hydroxypropionate
AAT: aspartate aminotransferase
CBB: Calvin Benson Bassham
DMA: Dimethylamine
HMA: High Microbial abundance
HMM: Hidden Markov Model
Lys: Lysine
MAGs: Metagenome assembled genomes
*mcrA*: methyl-coenzyme M reductase
MeA: Methylamine
MPN: Methylphosphonate
*mpnS*: methylphosphonate synthase
*mttB*: methyltransferase
QC: Quality control
rTCA: reductive Tricarboxylic Acid
SMP: Sponge microbiome project
SRBs: Sulfate reducing bacteria
TMA: Trimethylamine
TMAO: trimethylamine-N-oxide

## Availability of data and materials

All metatranscriptomes used in this study are publicly available under the following NCBI Bio project ID: XXXXX

## Acknowledgments

Livio Steindler is warmly thanked for assisting us as captain and for providing his sailing boat ‘Colpo de Fulmine II’ for the sponge sampling. We thank the ‘Area51 Diving School Trieste’ for supporting all the dive equipment in this research and assisting with the scientific dives. Cláudia Ferreira is acknowledged for the design of Figure 7. Dr. Stefan Green and Dr. Kevin J Kunstman from the Sequencing Core at the University of Illinois at Chicago (UIC) for advice on library preparation protocols and for sequencing. We thank Kelly Paglia and Christopher Brydges at the UC Davis metabolomic center for technical support with our sponge tissue analyses.

## Ethics Declarations

### Ethics approval and consent to participate

Not applicable.

### Consent for publication

Not applicable.

### Author Contributions

GAR: Project leadership, bioinformatic analyses, wrote the manuscript.

RBS: Laboratory work, data analysis.

AF: Experimental design of field work and sampling

RR: Experimental design of field work and sampling

MG: Experimental design of field work and sampling

GC: Experimental design of field work and sampling

AIG: Software development, data analysis.

LS: Project leadership, funding acquisition, analysis validation, field sampling, advised GAR, wrote the manuscript.

