## Supplementary material for "Bacterial aerobic methane cycling by the marine sponge-associated microbiome": Ramirez_SI_2022

**Conflict of interest**:

The authors declare no conflict of interest.

**Figure S1:** Amplicon Sequence Variants (ASVs) community composition and activity based on 16S rRNA genes amplified using as template either genomic DNA (gDNA, panel A) or complementary DNA (cDNA; synthesized from RNA; panel B) from the same four sponge specimens used in the metatranscriptomics libraries. Color codes represent the relative (A) community composition and (B) a proxy for the transcriptional activity of prokaryotic lineages classified at the Class-level of taxonomy. Each sponge specimen was sampled 24h apart at noon.

**Figure S2.** Abundance normalized activity survey for 37 dominant lineages, based on metatranscriptomic read mapping against MAGs, shown for all four sponge replicates sampled at noon 24h apart. Data is arranged based on phylogenomic relatedness and color-coded at the Phylum-level of taxonomy, in the *A. aerophoba*-associated microbiome. Relative activity is represented by circle size and also color-coded based on the Phylum-level association of each depicted lineage.

**Figure S3:** Complete KEGG-decoder pathway estimates heatmaps for all metatranscriptomic samples. Additional functional categories shown here and not in Figure 1 include: O_2_ cytochromes, hydrogen cycle, vitamins and transporters, secretion systems, and amino acid synthesis.

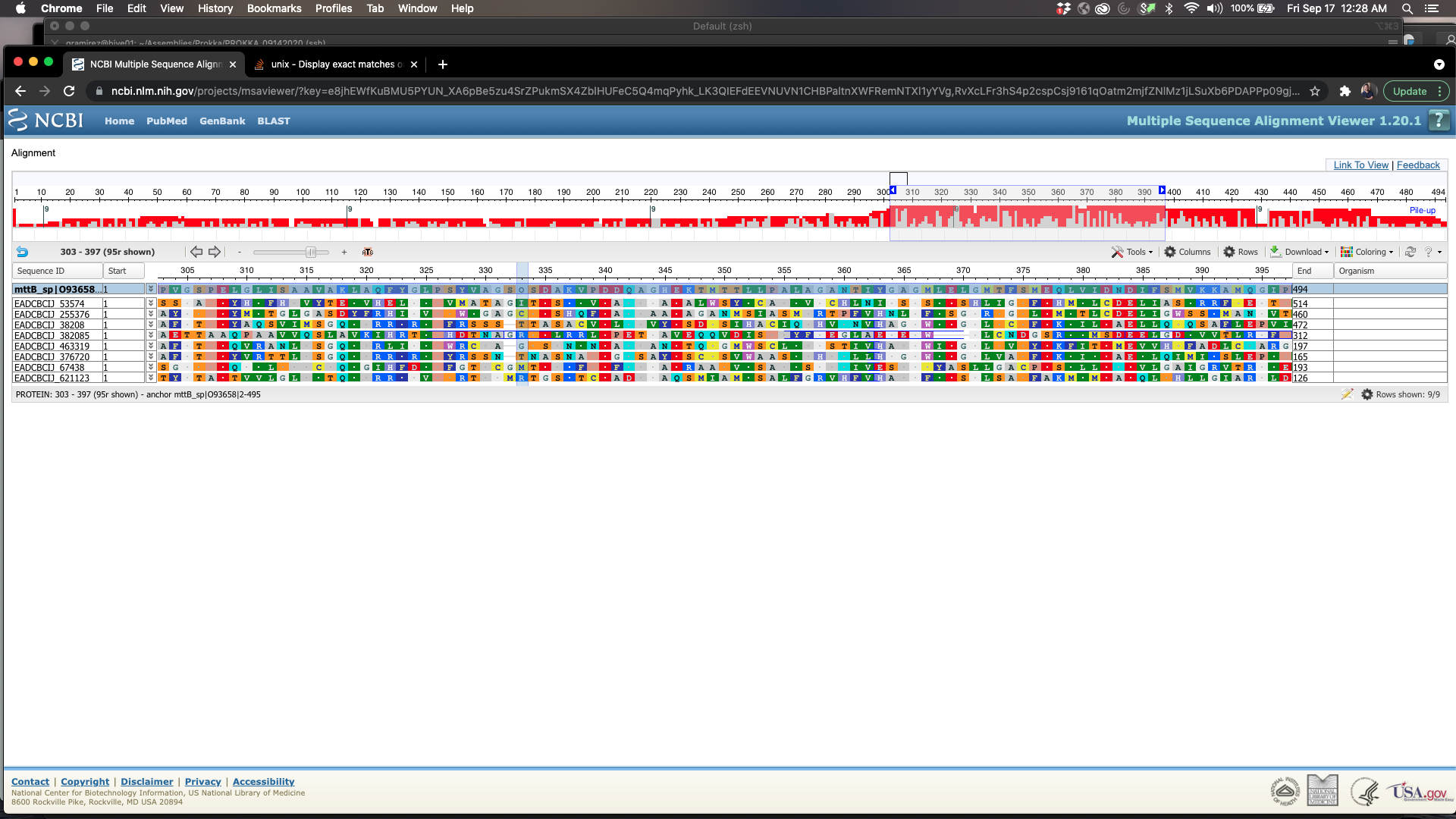

**Figure S4.** *mttB* gene transcript homologue alignment localized to a window between residues 303-440. The red column highlights residue 333, an expected non-conical pyrrolysine “O” residue found in all functional trimethylamine-corrinoid protein co-methyltransferases and absent in the sponge bacterial homologues.

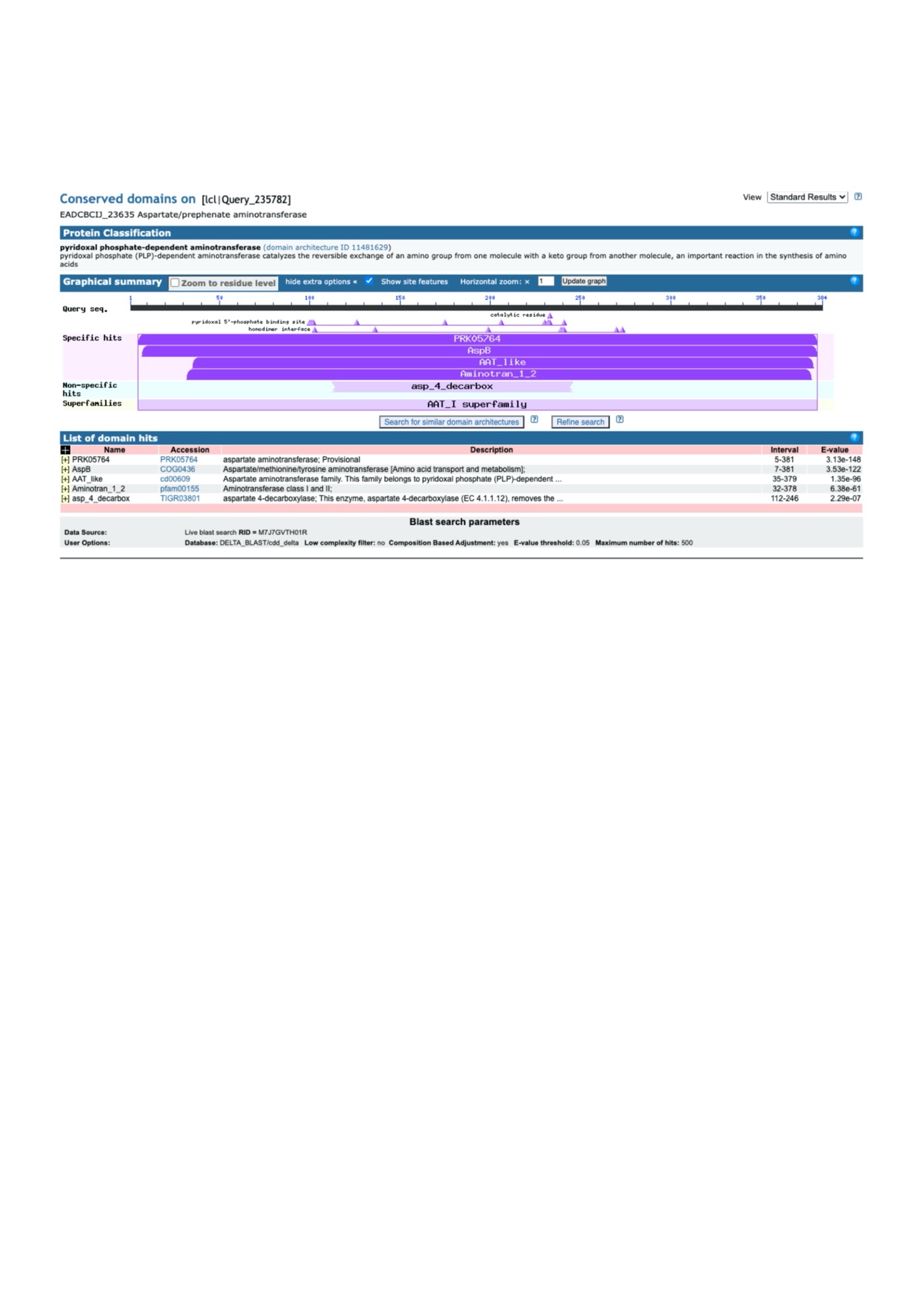

**Figure S5.** Delta-BLAST analysis for conserved functional motifs in proteins predicted from transcribed *aat* gene homologues.

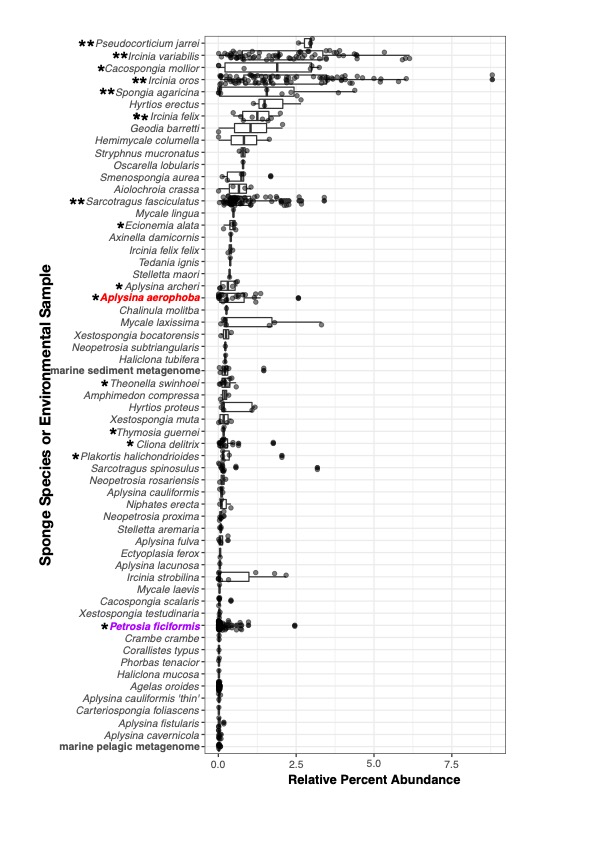

**Figure S6.** Relative percent abundance of 16S rRNA sequences with >98% sequence ID and 99% of query length to the *A. aerophoba* bin18 (Binatota) 16S rRNA gene sequence, summarized in quartile boxplots. Quartile box plots are arranged in descending median values of all observations from top to bottom. *Aplysina aerophoba* and *Petrosia ficiformis*, both sponges analyzed in this study, are highlighted in red and purple, respectively. Single and double asterisks depict sponge species with significantly (student t-test, P_val_ < 0.05) higher abundance of bin18-related sequences than in marine pelagic and marine sediment 16S rRNA datasets, respectively.

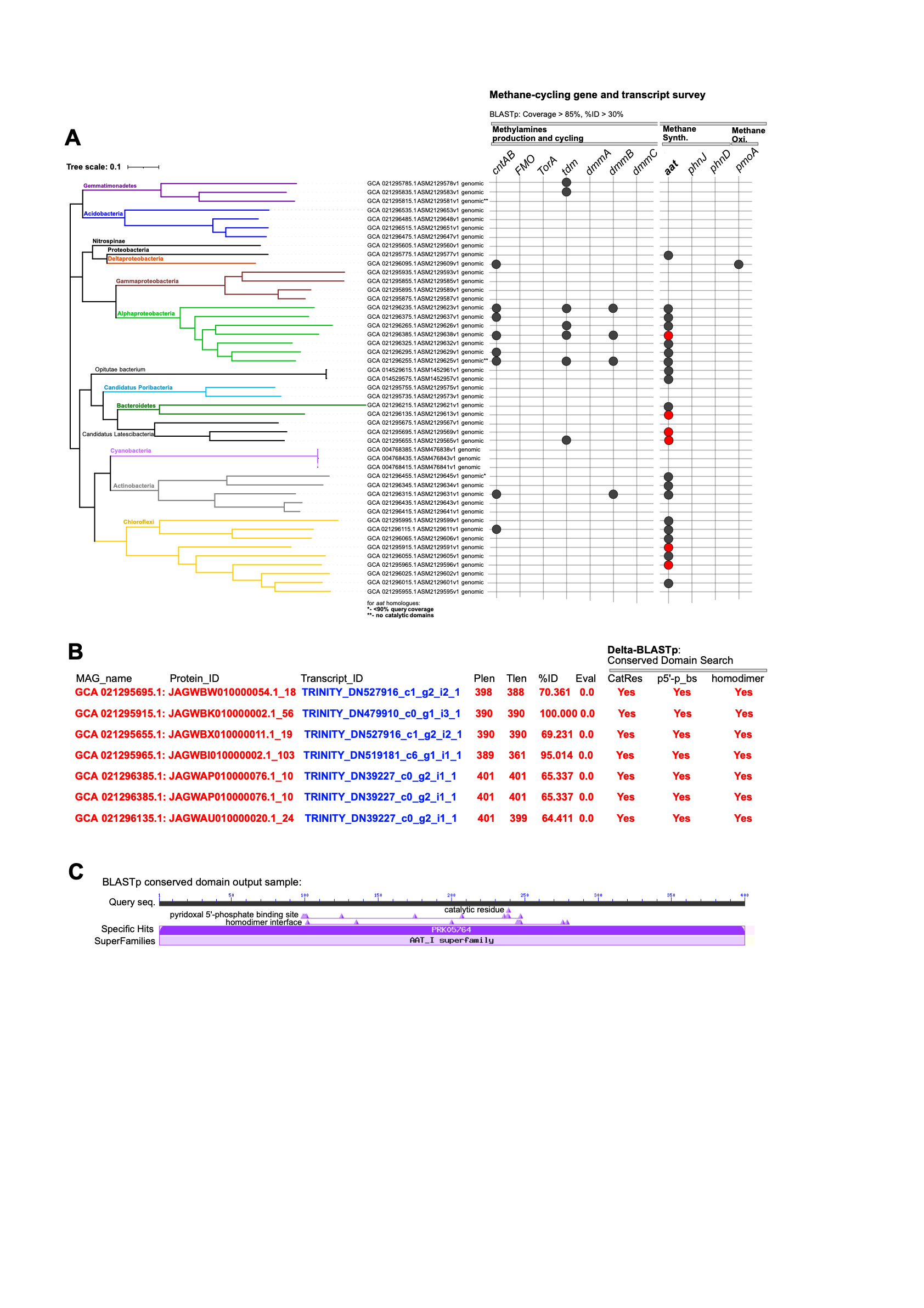

**Figure S7.** Meta-analysis of methane-cycling genes using public MAGs (arranged based on phylogenomic relatedness, (Burgsdorf et al. 2022) and metatranscriptomes from Mediterranean *P. ficiformis* sponges (Britstein et al. 2020). (A) Detected methane-related genes (black circles) and associated transcripts (red circles) for a collection of sponge symbiont MAGs. (B) Alignment metrics of metatranscriptome predicted (Transcript_ID) and MAG retrieved (Protein_ID) protein sequences identified as *aat* homologues including query (Tlen) and sequence length (Plen), %ID, Eval, and presence or absence of conserved domains. (C) Delta-BLASTp summaries of conserved domain position along a representative transcribed *aat* homologue sequence (all transcribed *aat* homologues shown in panel A contained identical conserved domain positions).

| **Library_ID** | **Library_ID2** | **Library Type** | **Raw_SeqCount** | **QC/Adapter/Ribodep** | **%Retained** |
| --- | --- | --- | --- | --- | --- |
| L36_S01_L001_004 | L36_L001_004 | Illumina PE | 27090089 | 14876193 | 54.91378415 |
| L38_S02_L001_004 | L38_L001_004 | Illumina PE | 27943100 | 13095154 | 46.86364076 |
| L39_S03_L001_004 | L39_L001_004 | Illumina PE | 24966863 | 12976736 | 51.97583693 |
| L40_S04_L001_004 | L40_L001_004 | Illumina PE | 27545844 | 11223495 | 40.74478531 |
| L66_S13_L001-004 | L66_L001_004 | Illumina PE | 31810768 | 17261353 | 54.26261007 |
| L68_S14_L001-004 | L68_L001_004 | Illumina PE | 27697230 | 12834651 | 46.33911406 |
| L69_S15_L001-004 | L69_L001_004 | Illumina PE | 25073626 | 11385626 | 45.40877335 |
| L70_S16_L001-004 | L70_L001_004 | Illumina PE | 28655209 | 11786638 | 41.1326192 |

**Table S1.** Metatranscriptomic read retention percentages following adapter trimming, interleaving, and rRNA library alignments for *in silico* rRNA depletion, prior to *de novo* assembly using rnaSPAdes.
